# Temporal and spatial profiling of ALKBH5 activity through NAIL-MS and compartmentalized RNA Isolation

**DOI:** 10.1101/2025.05.12.653425

**Authors:** Hagen Wesseling, Stefanie Kaiser

## Abstract

RNA modifications, especially m^6^A in human mRNA, are believed to be dynamically regulated through RNA writers and erasers. The key eraser of m^6^A is ALKBH5 with its function well proven in vitro, while in vivo evidence is lacking. Here, we set out to exploit nucleic acid isotope labelling coupled mass spectrometry (NAIL-MS) in a pulse chase set-up to study the in vivo function of ALKBH5 on human RNAs. For this we purified poly(A) from whole cell total RNA and found that, steady-state m^6^A levels and turnover dynamics were nearly identical between WT and ALKBH5 KO, despite clear evidence of robust RNA turnover within an 8-hour labeling period. To assess whether ALKBH5 might act in a compartment-specific manner, we employed an advanced subcellular fractionation strategy, allowing for the isolation of chromatin-associated, nucleoplasmic, and cytoplasmic RNA. These analyses confirmed that m^6^A accumulates during transcript maturation, with levels peaking in nuclear fractions and decreasing following export to the cytoplasm, supporting the thesis m^6^A is a dynamic modification. Notably, however, spatial and temporal profiles of m^6^A distribution and decay were unaffected by ALKBH5 KO. Even in chromatin-associated and nucleolar mRNA, where co-transcriptional modification and potential demethylation would be most plausible, m^6^A dynamics remained indistinguishable between WT and KO cells in the NAIL-MS context. We thus conclude that ALKBH5 has no major role in mRNA m^6^A demethylation in HEK 293T cells grown under optimal conditions.

## Introduction

N6-methyladenosine (m^6^A) represents the most abundant internal modification in mammalian messenger RNA (mRNA) and a dynamic regulation by a coordinated interplay of methyltransferases (“writers”) and demethylases (“erasers”) is claimed. The methyltransferase complex, composed of METTL3 and METTL14, catalyses the deposition of m^6^A on mRNA. Several m^6^A “reader” proteins recognize this modification and mediate downstream effects on mRNA metabolism, including translation, decay, stability, and nuclear export (as reviewed in (Schwartz 2016, Meyer and Jaffrey 2017)). The first m^6^A demethylase, FTO, was identified in 2011 (Jia, Fu et al. 2011), followed by the discovery of ALKBH5 as a second putative eraser in 2013 (Zheng, Dahl et al. 2013). Since then, the physiological relevance and mechanistic roles of these demethylases have remained subject of active debate. Subsequent research demonstrated that FTO preferentially demethylates m^6^Am at the 5′ cap rather than internal m^6^A sites under physiological conditions (Mauer, Luo et al. 2017). In contrast, the cellular context and conditions under which ALKBH5 exerts its proposed demethylase activity are still not fully understood. In murine models, ALKBH5 deficiency impairs fertility (Zheng, Dahl et al. 2013), and various studies have reported elevated ALKBH5 expression in cancer, correlating with reduced methylation of tumor suppressor-associated transcripts. In breast cancer cells, for instance, ALKBH5 promotes tumor progression by demethylating mRNAs encoding pluripotency factors, thereby stabilizing these transcripts and supporting the maintenance of cancer stem cells under hypoxic conditions (Zhang, Samanta et al. 2016). Similarly, acute myeloid leukemia has been associated with ALKBH5 overexpression, contributing to tumorigenesis (Shen, Sheng et al. 2020). However, recent findings complicate this picture. A study reported that neither ALKBH5 knock-out nor knock-down in cultured cells did significantly affect m^6^A levels, while only overexpression led to a measurable decrease (Stejskal, Rajecka et al. 2025). This observation contributes to the ongoing debate about the scope and biological significance of ALKBH5-mediated m^6^A demethylation, with current evidence suggesting limited activity under homeostatic conditions and a potentially enhanced role during cellular stress.

Given these discrepancies, it becomes increasingly necessary to revisit and critically assess the methodologies employed in studying the m^6^A-ALKBH5 axis. A critical step during sample preparation is the enrichment of mRNA. Total cellular RNA consist of approximately 80-85% ribosomal RNA (rRNA), 10% transfer RNA (tRNA) and only about 5% mRNA. mRNA enrichment is typically achieved through poly (A) selection, rRNA depletion (either by magnetic pulldown or RNase H based assays) or size based exclusion of unwanted RNAs. However, the purity of enriched transcripts must be carefully evaluated, as polyadenylated rRNA can be co-enriched during poly(A) selection (Slomovic, Laufer et al. 2006), and rRNA depletion protocols often fail to efficiently remove small rRNA species, as further illustrated in this study. Even after multiple rounds of poly(A) selection and rRNA pulldown, samples may still contain substantial amounts of rRNA (Legrand, Tuorto et al. 2017). Additionally, RNase H-based depletion methods can cause nonspecific RNA fragmentation (Wesseling, Krug et al. 2025). Following mRNA preparation, methylated RNA immunoprecipitation sequencing (MeRIP-seq) is the most widely used method for mapping m^6^A and assessing condition-dependent changes. It detects methylated regions as peaks in transcript coverage, comparing immunoprecipitated RNA to input RNA. However, the antibody used in MeRIP cannot reliably distinguish between m^6^A and m^6^Am (Linder, Grozhik et al. 2015) and MeRIP does not provide absolute quantification of methylation (stoichiometry). Moreover, MeRIP-based approaches suffer from low reproducibility due to high variability in signal, even among biological replicates from the same cell type (McIntyre, Gokhale et al. 2020). As an alternative, liquid chromatography–mass spectrometry (LC-MS) with single nucleosides allows for differentiation between m^6^A and m^6^Am but not the sequence the modification derived from. Each modified nucleoside has distinct physicochemical properties, resulting in specific retention times, and the *m/z* ratio provides a second characteristic feature for nucleoside identification. Absolute quantification of modified nucleosides is an additional benefit, if appropriate internal and external standards are employed (Kellner, Ochel et al. 2014). Nonetheless, LC-MS is particularly vulnerable to contamination from abundant non-mRNA species, especially rRNA and tRNA. Since mass spectrometers cannot distinguish whether a detected nucleoside originated from mRNA or another RNA species, contamination can significantly bias quantification.

Here, we present a temporally resolved analysis of mRNA methylation turnover using NAIL-MS technology (nucleic acid isotope labelling coupled mass spectrometry) (Heiss, Hagelskamp et al. 2021). Isotopically labelled nutrients were added to the cell culture medium, enabling distinction between pre-existing and newly synthesized m^6^A marks in total RNA and highly purified mRNA. With this approach we distinguish enzymatic demethylation through e.g. ALKBH5 from passive methylation loss due to transcript turnover or RNA decay. To assess whether m^6^A modification dynamics vary between nuclear subcompartments, we performed separate analyses of chromatin-associated and nucleoplasmic RNA fractions. mRNA integrity and purity were assessed via chip-based gel electrophoresis, and quantitative LC-MS/MS was used to simultaneously monitor marker modifications from tRNA and rRNA, serving as internal control for non-mRNA contamination. All in all, our data indicates that m^6^A is not actively turned over by ALKBH5 in our chosen HEK 293T cell model under homeostatic growth conditions.

## Results

### ALKBH5 knock out does not alter m^6^A modification levels in total cellular RNA

HEK 293T ALKBH5 knockout (KO) and corresponding wild-type (WT) cells were cultured until they reached 70% confluency. To assess m^6^A levels, total RNA was extracted and enzymatically digested into single nucleosides (Fig. 1A). These nucleosides were then quantitatively analysed *via* LC-QQQ-MS, using both internal and external standards (Ammann, Berg et al. 2023). To account for injection variability, m^6^A levels were normalised to guanosine (G) levels and the resulting value of m^6^A per 1000 G was plotted. As expected, comparison of WT and ALKBH5 KO cells revealed no significant difference in total cellular RNA m^6^A abundance (Fig 1B). Western blot analysis confirmed efficient ALKBH5 knockout (Fig. 1C and Supplemental Fig. S1A). Even after 23 passages, a second western blot (Supplemental Fig. S1B) confirmed the continued absence of ALKBH5, ruling out contamination by WT cells. Interestingly, RNA samples collected over multiple passages revealed a decreasing abundance of m^6^A abundance with increasing passage number (Supplemental Fig. S1C). Due to this observation, we were careful to perform all experiments with cells at 5-6 passages. For further analysis of m6A dynamics, we performed a pulse-chase NAIL-MS experiment. For this, cells were cultured to 70% confluency, and the addition of isotopically labelled nutrients (^15^N_5_-adenine, ^15^N_2_ ^13^C_5_-uridine and CD_3_-methionine) into the growth medium marks the starting time point (0 hours) (Fig. 1D, (Heiss, Borland et al. 2021)). After 8 hours continued growth, we could distinguish pre-existing transcripts from newly synthesized ones based on the mass shifts of all nucleosides. In the case of m^6^A, original nucleosides that are unlabelled show an *m/z* of 282. After incorporation of the new nutrients, two new isotopologues can be detected: the first is derived from original adenosine sites in original RNA that were methylated after addition of isotopically labelled nutrients and carry a CD_3_ mark instead of the CH_3_ mark. These “hybrid” m^6^As have an *m/z* of 285 and thus a +3 increase over the original m^6^A. The second isotopologue is the fully labelled (^15^N_5_ and D_3_) m^6^A nucleosides (called: news) with an *m/z* of 290. A complete list of the resulting masses is found in Supplemental Table S1.

**Figure 1:**
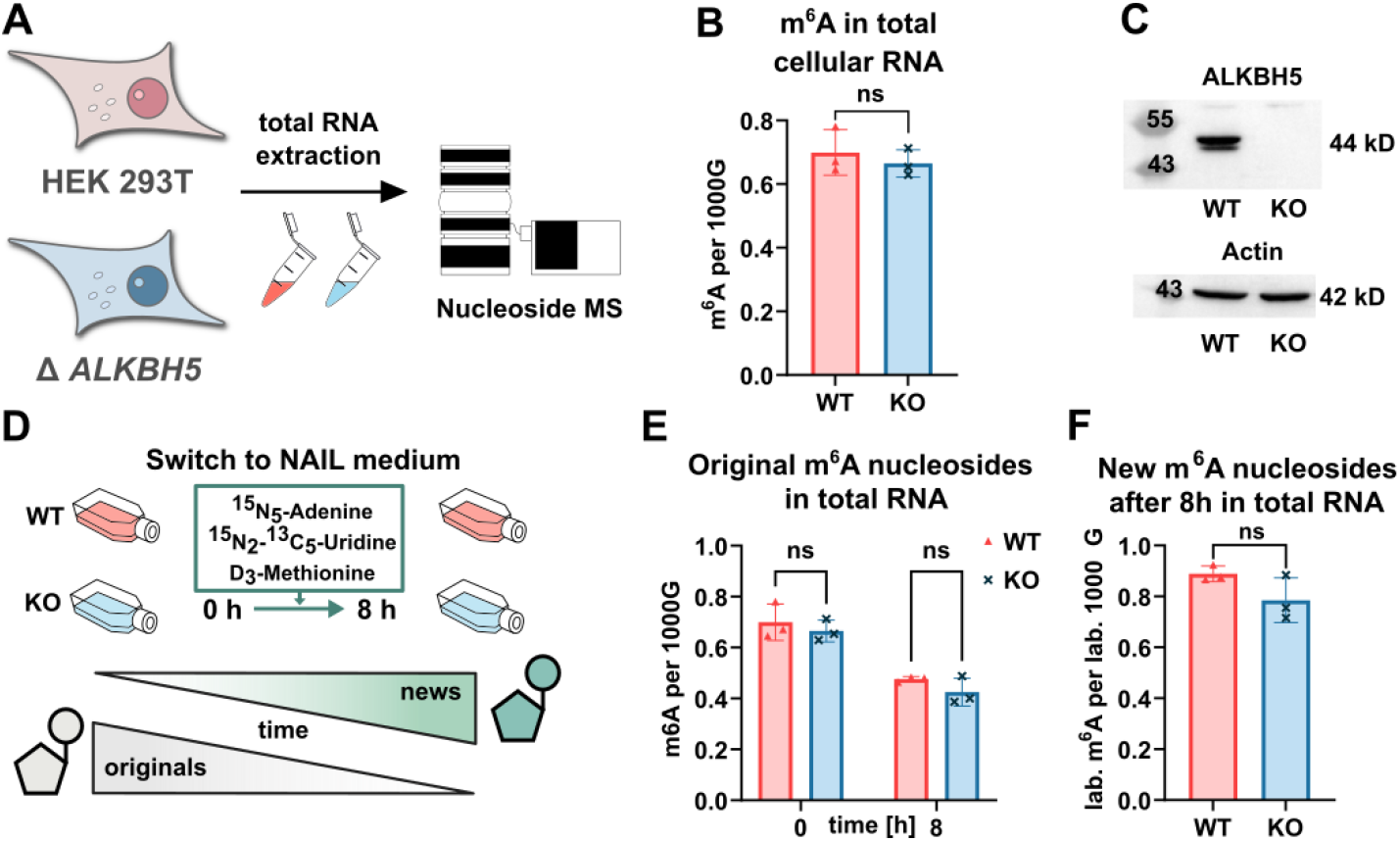
total RNA analysis of m^6^A abundance. **A:** total cellular RNA was harvested, digested to single nucleosides and analysed by LC/MS. **B:** Quantification results of m^6^A abundance normalized to 1000 guanosine (m^6^A per 1000 G) in HEK 293T and ALKBH5 deficient cells **C:** Western Blot with primary antibody for ALKBH5 (44 kD) and Actin (42 kD, control). Uncropped image and total Protein Staining shown in Supplemental Fig. S1A. **D:** Experimental setup of NAIL MS experiments to study temporal placement of RNA modifications **E & F:** Results of NAIL-MS study: fate of original m^6^A traced for 8h **(E)** and newly created m^6^A after 8h **(F)**. m^6^A concentrations in all experiments were normalized to 1000 Guanosines. Data was obtained from n=3 biological replicates. Statistical analysis by Welch’s t-test (ns = p > 0.05).

The stability of original m^6^A marks was monitored over an 8-hour period (Fig. 1E). The absolute abundance of original m^6^A drops both in WT and the ALKBH5 KO cells to a similar extent. The loss of original m^6^A independent of ALKBH5 indicates that the decrease is likely due to transcript degradation, especially of m^6^A-modified RNAs. The accuracy and precision of the NAIL-MS experiment is confirmed by the m^6^A abundance of newly transcribed m^6^A-containing RNA which is comparable to the original m^6^A abundance at experiment start (Fig. 1F). A steady-state modification profile of approximately 0.8 m^6^A per 1000 G appears to be maintained in total RNA of HEK 293T cells. Since m^6^A is not only the most abundant epitranscriptomic modification in mRNA but is also present in 18S rRNA (Ignatova, Stolz et al. 2020, Peng, Chen et al. 2022) and tRNA (*via* Dimroth re-arrangement from m^1^A to m^6^A (Macon and Wolfenden 1968)), we purified these RNAs by size exclusion chromatography (Supplemental Fig. S2A). The corresponding electropherograms are shown in Supplemental Fig. S2B. The small, but detectable, m^6^A amounts in tRNA potentially result from m^1^A re-arranging to m^6^A. No discernible differences in m^6^A were observed between wild-type and knockout samples (Supplemental Fig. S2C and D). These observations suggest that ALKBH5 does not substantially affect these RNA subtypes under the tested conditions. In a next step, we moved on to investigate m^6^A modifications specifically within mRNA.

### m^6^A abundance in whole cell derived poly(A) RNA is not affected by ALKBH5

To focus on mRNA-specific m^6^A modifications, we employed a combination of poly(A) enrichment and commercially available rRNA depletion using rcDNA (reverse complementary DNA) with magnetic bead-based pull-down. Cells from both WT and KO lines were harvested before and after an 8-hour NAIL-MS experiment. In the first replicate, a single round of poly(A) enrichment and rRNA depletion was performed, and RNA purity was assessed with chip gel electrophoresis. Supplemental Fig. S3A shows that residual 18S and 28S rRNA contamination remained, introducing potential biases in modified nucleoside quantification. To ensure complete removal of long rRNAs, a second round of poly(A) enrichment was performed in replicate two. The corresponding electropherograms (Supplemental Fig. S3B) demonstrate that while small RNAs (e.g., tRNA and 5/5.8S rRNA) were efficiently removed, additional rRNA depletion was still required. Doubling the amount of rcDNA probes successfully eliminated long rRNA contaminants, resulting in purified poly(A) RNA (Fig. 2A). Bioanalyzer data for all replicates are summarized in Supplemental Fig. S3C and D. Following purity validation, poly(A) RNA samples were enzymatically digested into nucleosides and analysed via LC-MS. As a marker for potential tRNA contamination, we quantified 5-methyluridine (m^5^U), a nucleoside predominantly found in tRNA (Richter, Plehn et al. 2022). The concentration of m^5^U was below the limit of quantification (LOQ) in the poly(A) RNA, indicating an abundance below 0.5% of potential tRNA contamination (calculation shown in Supplemental Fig. S3E). The respective rRNA marker, m^6,6^A, was also extremely low except for replicate one (Supplemental Fig. S2F). This was indeed expected from the gel electrophoresis (Supplemental Fig. S3A) and the replicate was excluded from further analysis. Data analysis of the remaining samples revealed a steady-state abundance of original m^6^A in poly(A) RNA of approximately 4 m^6^A per 1000 G, which decreased to ~1 m^6^A per 1000 G after 8 hours (Fig. 2B). The decrease in m^6^A is, again, independent of ALKBH5 which suggests a preferential degradation of m^6^A-containing poly(A) transcripts. Poly(A) RNA turnover is also visible from the new transcript ratio (NTR) which is the ratio of new (labelled) guanosine and original (unlabelled) guanosine (Fig. 2C). Interestingly, m^6^A abundance in new poly(A) RNA in the KO line is slightly lower than in WT (Fig. 2D), although we expected a higher abundance due to the missing demethylation activity of ALKBH5 in the KO. This effect is likely attributable to variations in cell density or mRNA isolation efficiency rather than ALKBH5 activity. A potential explanation for the non-detectable activity of ALKBH5 in the NAIL-MS context is that METTL3/14-mediated methylation and ALKBH5-mediated demethylation counteract each other with a demethylated site being masked by a (re-)methylated site. This dynamic equilibrium would result in an apparent constant m^6^A level. With NAIL-MS, these dynamics can be visualized by studying the hybrid m^6^As. Here, a demethylated and remethylated m^6^A would be detectable as a hybrid m^6^A signal due to the feeding with CD_3_-methionine. While we would expect a high abundance of “hybrid” turn-over m^6^A in WT, the abundance of post-methylated m_6_A should be lower in absence of ALKBH5. However, Figure 2E shows no significant difference, further supporting the conclusion that ALKBH5 does not significantly affect m^6^A levels in mature poly(A) RNA.

**Figure 2:**
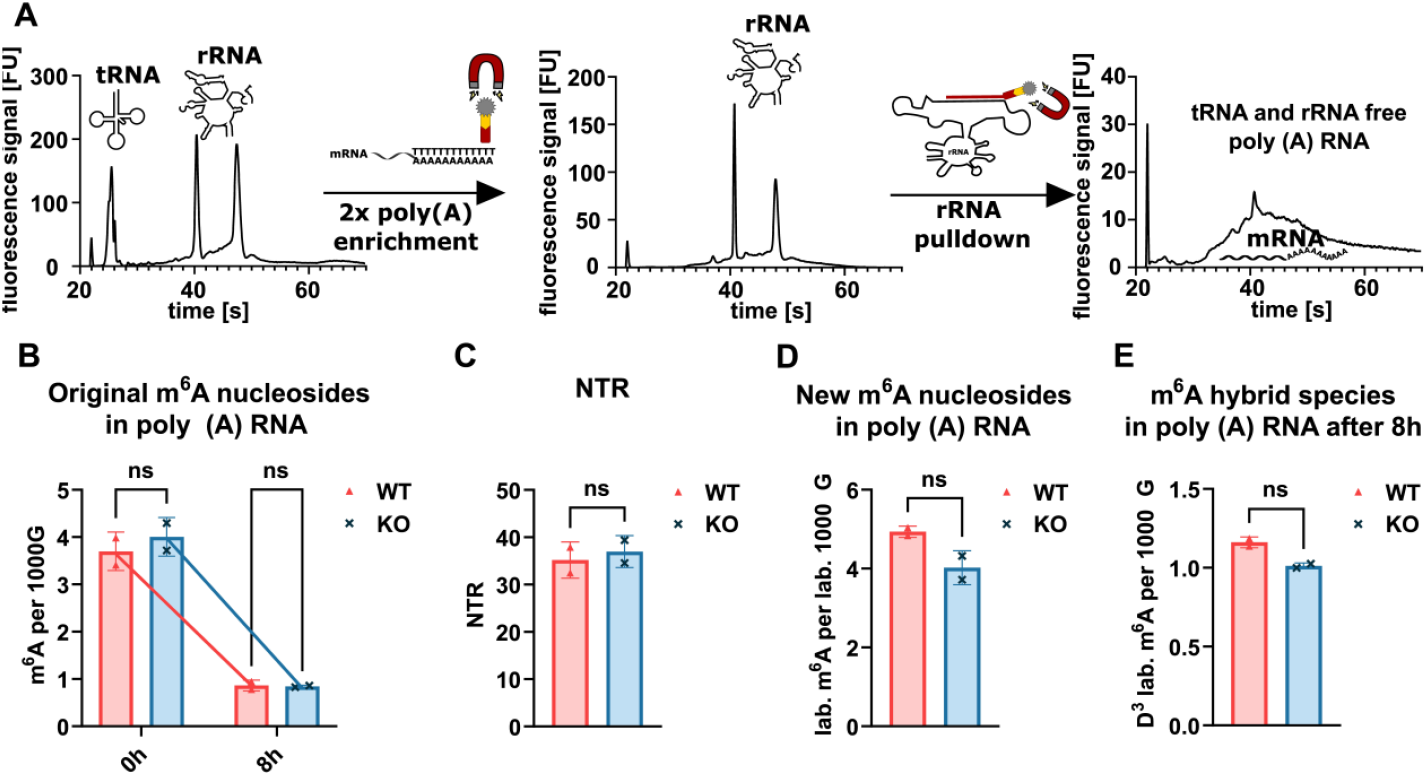
mRNA enrichment and analysis. **A:** Principle of rRNA depletion and poly (A) enrichment from total RNA samples. BioAnalyzer electropherograms indicate increasing mRNA purity. **B:** Fate of original m^6^A during 8h NAIL-MS experiment in HEK 293T and ALKBH5 deficient cells. **C:** Ratio of labelled Guanosine/all Guanosine isotopologues represents the new transcript ratio (NTR). **D:** NAIL-MS results of new generated m^6^A nucleosides (^15^N_5_ and D_3_) and **E:** hybrid m^6^A (D_3_) after 8h. m^6^A concentrations in all experiments were normalized to 1000 Guanosines. Data was obtained from n=2 biological replicates. Statistical analysis by Welch’s t-test (ns = p > 0.05).

### Subcellular fractionation allows isolation of RNA during its lifecycle

So far, we could not observe an effect of ALKBH5 on cellular RNA. However, our analysis has been limited to poly(A) RNA, which mainly contains mature mRNA from the cytosol, while ALKBH5 is reported to act in the nucleus (Zheng, Dahl et al. 2013). To investigate its role in nuclear RNA metabolism, we must study nuclear RNA in more detail. To do so, we adapted a cellular fractionation protocol (Conrad and Orom 2017) based on step-by-step lysis of the cytosolic and nucleic membranes allowing the isolation of chromatin-associated RNA (caRNA), nucleoplasmatic RNA (nucRNA) and cytosolic RNA (cytoRNA) (Fig 3A). To validate this approach and ensure that thus fractionated RNAs are available to NAIL-MS analysis, HEK 293T cells were cultured with isotopically labelled nutrients, and RNA from each subcellular compartment was harvested after 0, 2 and 8 hours. As RNA is transcribed from DNA located in the chromatin complex, we expect the highest NTR in caRNAs followed by nucRNAs. Afterwards RNA is exported from the nucleus to the cytoplasm, leading to a time-shifted increase in labelling and thus a lower NTR in cytoRNAs. The NTR was calculated and we found the expected differences in NTRs (Fig 3C). Gel electrophoresis of subcellular RNA fractions revealed that 45S rRNA precursors were present exclusively in caRNA, while mature 28S, 18S, and 5.8S rRNA appeared in nucleoplasmic and cytoplasmic fractions (Fig. 3B). tRNA levels were comparatively low in caRNA and nucRNA fractions, further validating fractionation efficiency. Following validation, we used this assay to analyse m^6^A modifications in both WT and KO cell lines. While most RNA modifications increased during transcript maturation from chromatin to cytoplasm (e.g. m^5^U in Fig 3E), m^6^A peaked in nuclear fractions and decreased upon nuclear export, consistent with a dynamic regulation process (Fig 3D). Despite previous reports that ALKBH5 demethylates nuclear RNA (Tang, Klukovich et al. 2018), our fractionation analysis did not reveal any ALKBH5-dependent effect on total nucRNA m^6^A abundance.

**Figure 3:**
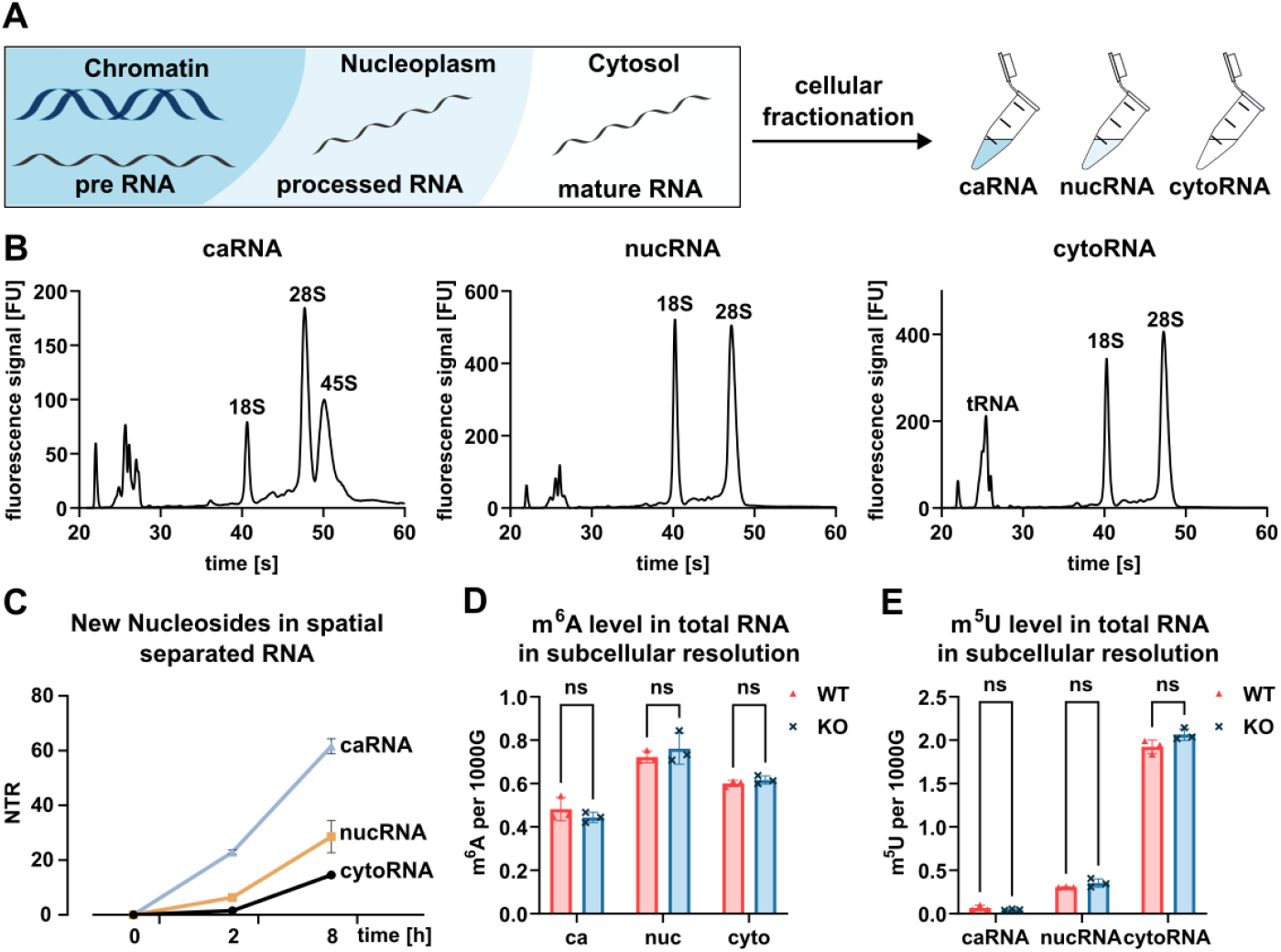
validation compartmentalised RNA isolation and results for total RNA. **A:** principle of cellular fractionation allowing isolation of chromatin-associated RNA (caRNA), nucleoplasmatic RNA (nucRNA) and cytosolic RNA (cytoRNA). **B:** electropherograms of isolated ca, nuc and cytoRNA marked with typical RNA subtypes (45S and tRNA). **C:** NAIL-MS experiment of HEK 293T cells subcompartimental fractionated at 0, 2 and 8h after addition of labelled nutrients. New transcript ratio (NTR) of each fraction was calculated as proof of concept. **D:** m^6^A in total RNA from subcellular compartments harvested from HEK 293T and ALKBH5 deficient cells. **E:** m^5^U levels in total RNA after subcompartimentalized fractionation of both cell lines. modification concentrations in all experiments were normalized to 1000 guanosines. Data was obtained from n=3 biological replicates. Statistical analysis by Welch’s t-test (ns = p > 0.05).

### ALKBH5 knockout shows no effect on m^6^A level in chromatin-associated and nucleoplasmic messenger RNAs

mRNA modifications occur both co- and post-transcriptionally. METTL3/14 (Louloupi, Ntini et al. 2018, Xu, Li et al. 2021) and ALKBH5 (Zheng, Dahl et al. 2013, Tang, Klukovich et al. 2018) have been reported to localize in nuclear speckles, where they regulate alternative splicing and nuclear export of mRNAs. To investigate m^6^A dynamics in chromatin-associated (ca) and nucleoplasmic (nuc) mRNAs, we established and validated an mRNA enrichment protocol independent of poly (A) selection, as pre-mRNAs do not possess quantitative poly (A) tails (Fig 4A).

**Figure 4:**
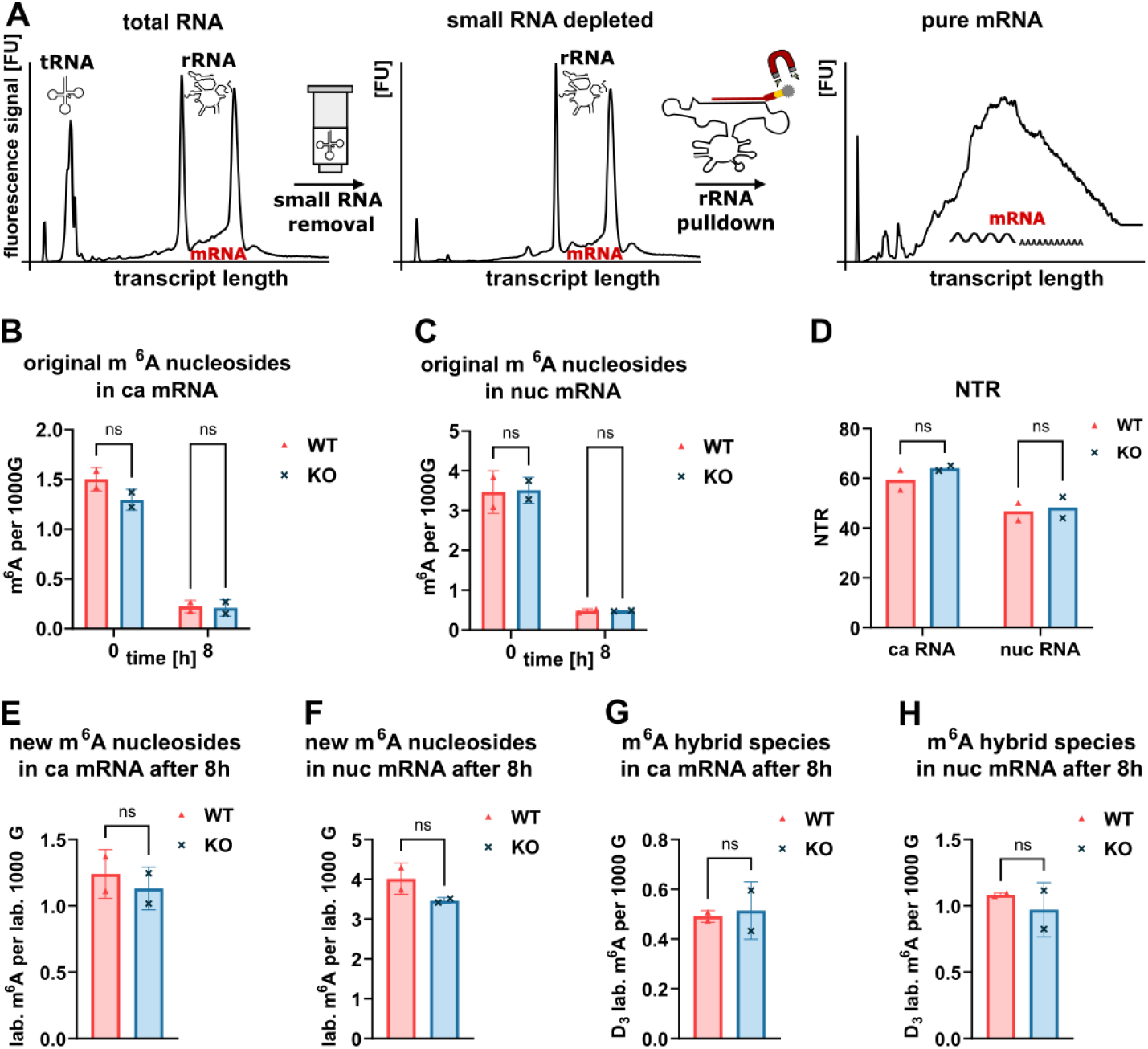
**A:** principle mRNA enrichment after subcellular fractionation and electropherogram results of mRNA after enrichment steps. **B and C:** Trace of mRNAs enriched from caRNA (**B**) and nucRNA **(C)** during 8h NAIL-MS experiment. **D:** Ratio of labelled (=new) Guanosine/all Guanosine isotopologues represents new transcript ratio (NTR) in ca and nuc mRNA fraction. **E and F:** New generated m^6^A nucleosides after 8h in NAIL medium shown for chromatin-associated mRNAs **(E)** and nucleoplasmatic mRNAs **(F). G and H:** NAIL-MS results for m^6^A hybrids after 8h NAIL context in ca mRNAs **(G)** and nuc mRNAs **(H)**. m^6^A concentrations in all experiments were normalized to 1000 Guanosines. Data was obtained from n=2 biological replicates. Statistical analysis by Welch’s t-test (ns = p > 0.05).

As a first step, cells were grown again for 8 hours in the described pulse-chase NAIL-MS setup and cells were harvested using the compartmentalized RNA isolation approach. A total of 10 *μ*g of ca and nuc RNA was obtained from each cell line in three biological replicates. For the first replicate, small RNAs (<200 nts, as specified by the manufacturer) were depleted from ca and nuc RNA using a commercial kit, followed by rRNA removal using twice the amount of rcDNA, as described in the previous section. Prior to LC-MS analysis, mRNA purity was assessed by chip-based gel electrophoresis. Unexpectedly, substantial amounts of small RNAs were still present (Supplemental Fig. S4A). In the commercial kit, ethanol is used for size-dependent precipitation of the RNA. Given that ethanol concentration determines the length of precipitated RNA during small RNA exclusion, we adjusted the ethanol concentration from 33% to 17% in a pilot experiment with ca total RNA. As shown in Supplemental Fig. S4B, lowering ethanol to 17% resulted in a marked reduction of small RNA contamination but also led to nearly complete loss of long RNAs. Consequently, we determined that an ethanol concentration of 23% provided an optimal balance between RNA yield and small RNA depletion efficiency. Small RNA depletion was repeated on the first replicate using 23% ethanol and now ~ 200 ng of mRNA were recovered from the 10 *μ*g input caRNA. Supplemental Fig. S4C shows the characteristic mRNA peak, confirming that these samples were suitable for LC-MS analysis. Replicates two and three were directly processed with 23% ethanol, requiring also two rounds of small RNA depletion to achieve efficient removal. Chip-based gel electrophoresis confirmed lower levels of residual small RNA contamination in these replicates (Supplemental Fig. S5A and B). Chromatin-associated and nucleoplasmic mRNAs from both cell lines at 0h and 8h time points were then subjected to LC-MS analysis. We assessed contamination with m^5^U (as tRNA marker) and m^6,6^A (as rRNA marker), as shown in Supplemental Fig. S5C-F. Replicate one exhibited persistently high levels of m^5^U and was therefore excluded from further analysis. While m^6,6^A was still detectable in most replicates, mRNA purity was maximized given the limited RNA input (100–200 ng total) required for LC-MS analysis of m^6^A and other modifications. At 0h, original ca-mRNAs contained 1.5 m^6^A per 1000 G in both cell lines, confirming the co-transcriptional deposition of m^6^A onto nascent transcripts (Fig. 4B). However, the majority of m^6^A modifications potentially occurs post-transcriptionally, as supported by the higher abundance of original m^6^A of 3.5 per 1000 G in nuc-mRNA (Fig. 4C) across both cell lines. After 8 hours, ca-mRNA retained 0.25 original m^6^A per 1000 G, corresponding to approximately 15% of its original levels at 0h. Due to the NAIL-MS context, the decrease is not caused by dilution with new caRNAs. A potential explanation might be a preferred release of m^6^A-modified RNAs from the chromatin complex und the nucleoplasm. In addition, this data indicates a high abundance of non-m^6^A modified RNAs in chromatin structures. This idea is supported by the NTR. Over the 8-hour period, 60% of caRNA transcripts were newly synthesized, indicating that 40% of the original transcripts remained chromatin-associated (Fig 4D). The discrepancy between 40% remaining transcripts and only 15% of the initial m^6^A levels suggests an m^6^A-dependent RNA transport or demethylation process. Regarding demethylation, both ALKBH5 knockout and wild-type cells exhibited the same pattern of original m^6^A loss and NTR. This indicates that ALKBH5 has no role in processing of caRNAs in these homeostatically grown HEK 293T cells.

In the nucleoplasmic fraction, original m^6^A levels also decreased from 3.5 to 0.5 m^6^A per 1000 G (~15%), with a NTR of ~45%. Similar to the chromatin-associated fraction, no evidence of ALKBH5 activity on nascent mRNA transcripts was observed in nucleoplasm. After 8 hours, newly synthesized transcripts contained 1.2 m^6^A per 1000 G in ca-mRNA in both cell lines (Fig. 4E). Similarly, nuc-mRNA exhibited increased methylation levels in newly transcribed mRNAs (Fig. 4F), yet no ALKBH5-dependent effect was detected. We previously hypothesized whether de- and remethylation mask each other which can be visualized by analysis of hybrid m^6^A. However, as shown in Fig. 4G and H, no ALKBH5-dependent effect on hybrid m^6^A levels was observed in either ca- or nuc-mRNA. Thus, we conclude that that ALKBH5 does not act on mRNA in unstressed HEK 293T cells.

## Discussion

This study set out to revisit the long-standing hypothesis that ALKBH5 functions as an active m^6^A demethylase in mRNA under basal conditions. Leveraging optimized mRNA enrichment workflows and subcellularly resolved NAIL-MS, we provide quantitative and subcellular-resolved insights into m^6^A dynamics in human cellular mRNA.

One of the key aspects we learned in this project is that absolute abundance of m^6^A in total RNA depends on the passage number. At this stage, we cannot explain why this is the case and whether this effect is exclusive for HEK 293T cells or a general effect. Thus, we recommend that the passage number should be noted alongside any data of an m^6^A-focused study and that the passage number should be kept low. Here, we have worked in cells in passage 5-6.

A second aspect we observed is that purification of mRNA, or better phrased poly(A) RNA if poly(A) enrichment techniques are employed, is extremely challenging and quality control in terms of size analysis is needed to assess the presence of rRNA and small RNA. While we evaluated multiple enrichment strategies, we found that there is not one protocol to fit all samples types, but rather the method must be tailored to specific sample types.

Finally, we want to emphasize the utility of deconvolution and reduction of sample complexity to gain insight into complex biological questions. By combining the temporal resolution afforded by NAIL-MS with spatial resolution through subcellular fractionation, we provide a solid foundation to truly study m^6^A dynamics. Although the data in this manuscript is only a small step forward to a global understanding of m^6^A dynamics, our data challenges parts of the model proposed in 2013 (Zheng, Dahl et al. 2013), namely that ALKBH5 acts as a major m^6^A demethylase *in vivo*. Instead, our results support recent findings (Stejskal, Rajecka et al. 2025) reporting minimal impact of ALKBH5 depletion on global m^6^A levels. While ALKBH5 may play important roles under specific physiological or pathological contexts, such as hypoxia (Zhang, Samanta et al. 2016, Chao, Shang et al. 2020), cancer (Liang, Zhu et al. 2022), hematopoiesis (Gao, Zimmer et al. 2023), or differentiation (Han, Xu et al. 2021), its role in maintaining m^6^A homeostasis under basal conditions in HEK293T cells appears negligible.

## Materials & methods

### Chemicals and reagents

All salts, reagents and media were obtained from Sigma-Aldrich (Munich, Germany) unless stated otherwise. The isotopically labelled compounds ^13^C_5_,^15^N_2_-Uridine were obtained from Silantes (Munich, Germany) and ^15^N_5_-Adenine was obtained from Cambridge Isotope Laboratories (Tewksbury,MA, USA). All solutions and buffers were made with water produced in-house using a MilliQ EQ 7000 system (Merck Millipore, Burlington, MA, USA). The nucleosides adenosine (A), cytidine (C), guanosine (G) and uridine (U), were obtained from Sigma-Aldrich. N6-methyladenosine (m^6^A) and 5-methyluridine (m^5^U) were obtained from Carbosynth (Newbury, UK), while 6,6-dimethyladenosine (m^6,6^A) was obtained from Alfa Chemistry (Zhengzhou City, China).

### Cell culture

HEK 293T and ALKBH5 KO (ab266762, abcam, Cambridge, UK) cells were cultured at 37 °C in an atmosphere containing 10% CO_2_ for pH regulation, provided by a CellXpert C170i incubator (Eppendorf, Hamburg, Germany). For standard culture, DMEM D6546 (high glucose) was used, supplemented with 10% fetal bovine serum (FBS), 0.584 g/l L-glutamine, and 8 μg/l queuine. Cells were passaged at a 1:5 ratio every 2 days to prevent overgrowth. To avoid contamination, only sterile consumables were used, and all procedures were performed under a laminar flow hood (HeraSafe 2025, Thermo Fisher Scientific, Waltham, MA, USA). For nucleic acid isotope labeling experiments, DMEM D0422 (lacking cysteine and methionine) was used. The medium was supplemented with 10% dialyzed FBS, 0.584 g/L L-glutamine, 0.063 g/L cysteine, 0.03 g/L labelled methionine, 0.05 g/L labelled uridine, 0.015 g/L labelled adenine, and 8 μg/L queuine.

### RNA isolation

For RNA isolation, the culture medium was aspirated, and 1 mL of TRI Reagent (R2050, Zymo, Irvine, CA, USA) was added to 6 × 106 cells according to the manufacturer’s instructions. After vigorous mixing to lyse the cells, 200 μL of chloroform were added and mixed thoroughly. Samples were incubated for 10 minutes at room temperature, followed by centrifugation at 12,000 × g for 10 minutes. The aqueous phase (550 μL) was transferred to a new tube, mixed with 550 μL of isopropanol (102781, Merck, Darmstadt, Germany), and incubated overnight at −20 °C. RNA was pelleted by centrifugation at 12,000 × g for 60 minutes at 4 °C, washed with 200 μL of 70% ethanol for 15 minutes (12,000 × g, 4 °C), and resuspended in 50 μL of ultrapure water. Total RNA concentrations were measured using an N60 nanophotometer (Implen GmbH, Munich, Germany).

### Subcellular fractionation

Cellular fractionation of cultured cells was performed based on a previously published protocol (Conrad and Orom 2017) with minor modifications. All sample handling steps were carried out on ice, and all buffers were supplemented with 20 U/mL SUPERase RNase Inhibitor (AM2694, Thermo Fisher Scientific, Waltham, MA, USA). A total of 6 × 10^6^ HEK 293T cells were cultured in T25 flasks and detached by trypsinization. After adding 4 mL of cold culture medium, the cell suspension was centrifuged at 300 × g for 3 minutes. The resulting pellet was resuspended in 1 mL PBS and transferred to 1.5 mL reaction tubes. Following a second centrifugation at 300 × g for 3 minutes, 400 *μ*L of lysis buffer (10 mM Tris, pH 7.4 (4855.2, Roth, Karlsruhe, Germany), 150 mM NaCl (0601.1, Roth, Karlsruhe, Germany), and 0.15% Igepal CA-630 (J61055, Thermo Fisher Scientific, Waltham, MA, USA)) were added. After resuspending the pellet, samples were incubated on ice for 5 minutes. The lysate was then carefully layered onto 1 mL of sucrose buffer (24% sucrose, 10 mM Tris, pH 7.4, and 150 mM NaCl) in a fresh tube. The cytoplasmic fraction (supernatant) was collected and after high speed centrifugation (15,000 x g, 1 min, 4°C) transferred in a new reaction tube. The remaining pellet (nuclear fraction) was washed gently with PBS-EDTA, followed by the addition of 250 *μ*L glycerol buffer (20 mM Tris, pH 7.4, 75 mM NaCl, 0.5 mM EDTA, 50% glycerol) and nuclear lysis buffer (10 mM Tris, pH 7.4, 1 M Urea (3941.1, Roth), 0.3 M NaCl, 0.2 mM EDTA, 1% Igepal CA-630). After a 2-minute incubation on ice, samples were centrifuged (13,000 x g, 2 min, 4°C) resulting in a chromatin pellet and a supernatant containing nucleoplasmic fraction. The chromatin pellet was washed with PBS-EDTA and subsequently dissolved in 1 mL TRI Reagent by vortexing. For RNA isolation, 1 mL of TRI Reagent was added per 0.2 mL of collected cytoplasmic or nucleoplasmic lysate. After vortexing, 200 *μ*L chloroform were added to each tube, followed by a 10-minute incubation at room temperature. Samples were centrifuged at 12,000 x g for 10 minutes. The aqueous phase (550 *μ*L) was transferred to a new tube, mixed with 550 *μ*L isopropanol (102781, Merck, Darmstadt, Germany), and incubated overnight at −20 °C. RNA was pelleted by centrifugation at 12,000 × g for 60 minutes at 4 °C, washed twice with 200 *μ*L of 70% ethanol (15 minutes each, 12,000 × g, 4 °C), and finally resuspended in 30 *μ*L of ultrapure water.

### mRNA enrichment from total cellular RNA

To isolate poly(A) RNA the magnetic mRNA Isolation Kit (S1550S, NEB, Ipswitch, USA) was used according to the manufacturers protocol, with following minor changes: 50 *μ*g total RNA was added to 500 *μ*L Bind/Lysis Buffer in Step 2. The volume of the Elution buffer was decreased to 46 *μ*L. To deplete rRNA leftovers the eluted poly(A) RNA was treated with an RiboMinus Eukaryote Kit v2 (A15020, Thermo Fisher Scientific, Waltham, MA, USA). Here, the amount of the rcDNA (eukaryote probe mix v2) was increased to 8 *μ*L. After proceeding, mRNA was washed by ethanol precipitation and recovered in 10 *μ*L ultrapure water.

### mRNA enrichment from chromatin-associated and nuclear RNA

Small RNAs (< 200 nts) were removed using the RNA Clean & Concentrator Kit (Zymo Research, Irivine, CA, USA) with 5-10 *μ*g of total RNA as input material. Ethanol concentration was decreased to 23 %. To deplete rRNA leftovers the eluted poly(A) RNA was treated with an RiboMinus Eukaryote Kit v2 (A15020, Thermo Fisher Scientific, Waltham, MA, USA). Here, the amount of the rcDNA (eukaryote probe mix v2) was increased to 8 *μ*L. After proceeding, mRNA was washed by ethanol precipitation and recovered in 10 *μ*L ultrapure water. Remaining small RNAs were then removed by again using the RNA & Concentrator Kit (Zymo Research, Irvine, CA, USA).

### size exclusion chromatography

tRNA and rRNA were purified by size exclusion chromatography using an Agilent 1100 Series system (Agilent Technologies, Santa Clara, CA, USA) and a BioSEC Advance 300 Å column (2.7 μm, 7.8 × 300 mm; PL1180-5301, Agilent Technologies). The chromatography was performed with 0.1 M ammonium acetate as the mobile phase at an isocratic flow rate of 1 ml/min. The column temperature was maintained at 40 °C. Approximately 10 μg of total RNA was loaded per run. tRNA and rRNA were separated by size and collected based on retention time using an automated fraction collector. The collected sample was concentrated to ~50 μL using a Savant SpeedVac SPD120 (Thermo Fisher Scientific, Waltham, MA, USA). For RNA precipitation, 0.1 volumes of 5 M ammonium acetate and 2.5 volumes of ice-cold ethanol were added. Samples were incubated overnight at −20 °C, followed by centrifugation at 12,000 x g for 60 minutes at 4 °C.

### Digestion prior LC-MS

500 ng total RNA (or 500 ng rRNA and 300 ng tRNA) of each sample were hydrolysed to single nucleosides in a volume of 35 *μ*L by using 2 U alkaline phosphatase, 0.2 U phosphodiesterase I and 2 U benzonase in Tris (pH 8, 5 mM) and MgCl_2_ (1 mM) containing buffer. Furthermore, 0.5 *μ*g tetrahydrouridine, 1*μ*M butylated hydroxytoluene and 0.1 *μ*g pentostatin were added to avoid deamination and oxidation of the nucleosides. For mRNA analysis, the volume of the digest was decreased to 20 *μ*L by adding 0.8 U alkaline phosphatase and Benzonase and 0.08 U phosphodiesterase 1 to 50-200 ng enriched mRNA. Additionaly, digestion mix contains 0.4 *μ*g Pentostatin, 2 *μ*g THU and 4 *μ*M BHT. After incubation for 2 h at 37 °C, 15 *μ*L, in case of mRNA analysis: 5 *μ*L, of LC-MS buffer (5 mM NH_4_OAc, brought to pH 5.3 with glacial acetic acid) was added. For LC-MS/MS analysis, 10 *μ*L of each total RNA sample, respectively 15 *μ*L of each mRNA sample, were injected and 1 *μ*L stable isotope labelled internal standard (SILIS) spiked into.

### Nucleoside mass spectrometry

For quantitative mass spectrometry an Agilent 1290 Infinity II combined with an Agilent Technologies G6470A QQQ system and electrospray ionization was used. For separation of Nucleosides a Fusion-RP column at 35 °C and a flow rate of 0.35 ml/min was used. Buffer A was LC-MS buffer, buffer B was Ultra LC-MS grade acetonitrile. The gradient started at 100% solvent A for 1 min, followed by an increase to 10% solvent B over 4 min. From 5 to 7 min, solvent B was increased to 40% and maintained for 1 min before returning to 100% solvent A in 0.5 min and a 2.5 min re-equilibration period. The instrument was operated in dynamic MRM mode. Total RNA analysis was performed with an EMV of 200, increased to 500 for mRNA analysis.

### Quantification of Nucleosides

For calibration, synthetic nucleosides were weighed and dissolved in water to a stock concentration of 1–10 mM. The calibration solutions ranged from 0.025 to 100 pmol for each canonical nucleoside and from 0.00125 pmol to 5 pmol for each modified nucleoside. Each calibration was spiked with 1 *μ*L SILIS. Data was analysed by the quantitative and qualitative MassHunter Software from Agilent. The areas of the MRM signals were integrated for each modification and their isotopologues. All samples and calibration signals were quantified using SILIS, as described earlier (Heiss, Borland et al. 2021). The absolute amounts of the modifications were normalized to the absolute amounts of Guanosine multiplied by 1000. In the case of the pulse-chase experiment, the different isotopomers were referenced to their respective labelled canonicals, so that original modifications were referenced to original canonicals and new modifications were referenced to new canonicals.

### Protein Extraction and Western Blotting

Cells were harvested using 1-minute trypsinization in a T25 flask, followed by the addition of 2 mL cell culture medium to neutralize trypsin. Cells were centrifuged at 300 × g for 3 minutes, washed once with 5 mL PBS, and resuspended in 100 *μ*L of RIPA buffer. Cells were incubated on ice for 30 minutes, during which they were pipetted up and down every 5 minutes to ensure thorough lysis. The samples were subsequently centrifuged at 13,000 × g for 15 minutes at 4°C. The resulting supernatant, containing total cellular protein, was collected. For both WT and ALKBH5 KO samples, 20 *μ*g of total protein were prepared for SDS-PAGE. Proteins were mixed with an equal volume of 2× Lämmli buffer and denatured by heating at 95°C for 5 minutes. Electrophoresis was performed via SDS-PAGE for 30 minutes (40 mA). Proteins were transferred to a PVDF membrane (abcam, Cambridge, UK) for 30 minutes. Post-transfer, the membrane was divided into two halves. One half was subjected to No-Stain total protein staining (A44449, Invitrogen, Carlsbad, CA, USA) by incubating in 20 mL of No-Stain solution (prepared by combining 19 mL H_2_O, 1 mL 20X buffer, and 40 *μ*L of both No-Stain reagents) for 10 minutes on a rolling platform, followed by two washes with water. This half was then blocked with 10 mL 5% BSA (8076.5, Roth, Karlsruhe, Germany) in TBST buffer for 1 hour at room temperature, after which 5 *μ*L of primary antibody against ALKBH5 (rabbit, HPA007196, sigma resp. Atlas Antibodies) was added to the blocking solution for overnight incubation. After washing with TBST buffer, HRP-linked Anti-rabbit-antibody (ab6721, abcam, Cambridge, UK) was added 1 hour prior to imaging. The second membrane half was incubated for 1 hour at room temperature on a rolling platform with an actin antibody (A3854, sigma) directly conjugated to peroxidase. Clarity Western ECL substrate was subsequently applied, and imaged using a Bio-Rad Chemidoc XRS+ Molecular Imager.

## Supplemental materials

Supplemental material is available for this article.

## Acknowledgements

This work was supported by the Deutsche Forschungsgemeinschaft [325871075-SFB 1309 to S.K.].

